# Population dynamics of epithelial-mesenchymal heterogeneity in cancer cells

**DOI:** 10.1101/2022.01.21.475826

**Authors:** Paras Jain, Sugandha Bhatia, Erik W Thompson, Mohit Kumar Jolly

**Affiliations:** Centre for BioSystems Science and Engineering, Indian Institute of Science, Bangalore, India; Queensland University of Technology, School of Biomedical Sciences, Faculty of Health, and Translational Research Institute, Brisbane, Australia; The University of Queensland Diamantina Institute, The University of Queensland, Brisbane, QLD, Australia

**Keywords:** Asymmetric Cell Division, Epithelial-Mesenchymal Heterogeneity, Epithelial-Mesenchymal Plasticity, Population Dynamics

## Abstract

Phenotypic heterogeneity is a hallmark of aggressive cancer behaviour and a clinical challenge. Despite much characterisation of this heterogeneity at a multi-omics level in many cancers, we have a limited understanding of how this heterogeneity emerges spontaneously in an isogenic cell population. Some longitudinal observations of dynamics in epithelial-mesenchymal heterogeneity, a canonical example of phenotypic heterogeneity, have offered us opportunities to quantify the rates of phenotypic switching that may drive such heterogeneity. Here, we offer a mathematical modeling framework that explains the salient features of population dynamics noted in PMC42-LA cells: a) predominance of EpCAM^high^ subpopulation, b) re-establishment of parental distributions from the EpCAM^high^ and EpCAM^low^ subpopulations, and c) enhanced heterogeneity in clonal populations established from individual cells. Our framework proposes that fluctuations or noise in content duplication and partitioning of SNAIL – an EMT-inducing transcription factor – during cell division can explain spontaneous phenotypic switching and consequent dynamic heterogeneity in PMC42-LA cells observed experimentally at both single-cell and bulk level analysis. Together, we propose that asymmetric cell division can be a potential mechanism for phenotypic heterogeneity.

## Introduction

Intra-tumor heterogeneity is a major roadblock that thwarts multiple therapeutic approaches in the clinic [1]. It has earlier been largely thought of as existing at a genomic level, i.e. co-existence of many sub-clonal populations. Single-cell genomic analysis has helped construct the lineage trees mirroring clonal evolution [2]. However, recent preclinical (*in silico, in vitro, in vivo*) and clinical observations have emphasized that besides genetic heterogeneity, tumors exhibit substantial non-genetic heterogeneity as well, often referred to as phenotypic heterogeneity [3–5]. Non-genetic heterogeneity can facilitate ‘bet hedging’ in a cancer cell population, thus enhancing its fitness under stressed conditions (immune attack, targeted therapy etc.) and enabling survival of subpopulations that can eventually drive tumor relapse and/or metastasis [6,7]. Therefore, identifying the mechanisms underlying non-genetic heterogeneity are of fundamental importance.

A canonical example of intra-tumor phenotypic heterogeneity is along the epithelial-mesenchymal axis. Epithelial-Mesenchymal Transition (EMT) and its reverse Mesenchymal-Epithelial Transition (MET) were initially considered as binary processes, but recent investigations across carcinomas, especially those at a single-cell level, have demonstrated that cancer cells can display many hybrid epithelial/mesenchymal (E/M) phenotypes *in vitro* and *in vivo*, as well as in patient samples [8–16]. Depending upon the combination of markers used in a specific study, cancer cells can be classified into two or more phenotypes – Epithelial (E), Mesenchymal (M) and the hybrid E/M one(s) [17]. However, most studies focus on a static snapshot of E-M heterogeneity, with little longitudinal data that can help unravel the set of underlying mechanisms initiating and sustaining this heterogeneity.

A few investigations into the population dynamics of E-M heterogeneity has revealed that when these phenotypically diverse subpopulations of cells are sorted by FACS (Fluorescent activated cell sorting) and cultured independently, over time, they can often give rise to other phenotypes in the parental population. These observations are reminiscent of stochastic cell-state transitions seen among cancer stem cells (CSCs) and non-CSCs [18]. For instance, any of the three (E, M, hybrid E/M) subpopulations isolated and cultured from prostate tumor cells (*PKV* cell line) could gave rise to other subpopulations in different proportions within two weeks *in vitro* [8]. Similarly, *in vivo*, subcutaneous transplantation of different SCC tumor subpopulations with varied EMT status led to a sustained co-existence of diverse phenotypes in the corresponding tumors [9]. These trends indicated the role of bidirectional phenotypic plasticity in promoting the emergence of E-M heterogeneity.

The phenotypic distribution of a cell population can vary across cell lines and single-cell clones generated from a cell line. For example, in a study across six different breast cancer cell lines, while four of them were largely homogenous in terms of relative levels of EpCAM (Epithelial Cell Adhesion Molecule – a common epithelial marker), two of them - HCC38 and HCC1143 - had a 90:10 and 99:1 ratio of EpCAM^high^ to EpCAM^low^ cells respectively [19]. Similarly, the PMC42-LA cell line comprised 80% EpCAM^high^ cells and 20% EpCAM^low^ [20], with the latter showing canonical mesenchymal morphological (spindle-shaped) and molecular (higher levels of EMT-transcription factors SNAIL, SLUG, ZEB1 and mesenchymal markers VIM and FN1) traits. When these two subpopulations were segregated and cultured separately, they returned to a 80:20 parental population distribution within 8 weeks. However, the single-cell clones established from PMC42-LA showed a more diverse phenotypic distribution in terms of ratios of EpCAM^high^ to EpCAM^low^ cells. Importantly, these different clones had varied migratory, invasive, tumor-initiating and drug resistance features, indicating that the ratio of cells in different phenotypes can influence the overall ‘fitness’ of the population for invasion-metastasis cascade. Similar molecular and functional diversity for single-cell clones was reported in another breast cancer cell line SUM149PT [21]. However, how these different subpopulation ratios are achieved and maintained remains elusive.

Here, we show, using a mathematical modelling approach, that in a cell population carrying the EMT regulatory network (miR-200/ZEB/SNAIL) [22], noise or fluctuations in processes of content duplication and partitioning of biomolecules can drive asymmetric cell division and can explain the observations for PMC42-LA system. We consider the influence of these fluctuations on the inherited levels of EMT-transcription factor SNAIL by the two daughter cells; The extent of these fluctuations has been assumed to be proportional to SNAIL levels of the dividing parent cell. As SNAIL regulates the levels of ZEB and miR-200 in a cell – that collectively define its EMT status [22,23] – we can recapitulate the spontaneous phenotypic switching among subpopulations with varied EMT status. Our model simulations can explain the observations in PMC42-LA cells – a) the dominance of EpCAM^high^ subpopulation over EpCAM^low^ subpopulation, and b) heterogeneity in EpCAM profile in single-cell clones. Thus, our results propose a possible mechanism that may underlie how non-genetic heterogeneity is generated in an isogenic cancer population.

## Results

### Dominance of epithelial cells in the population over time irrespective of initial distribution

Here, we have developed a population dynamics framework to explain the emergence of epithelial-mesenchymal heterogeneity in a given population, and contribution of spontaneous state switching in enabling this heterogeneity (**Fig 1A**). Specifically, we consider phenotypic switching to occur during cell division (**Fig 1B**), where two factors can contribute to a daughter cell having a phenotype different than its parent cell: noise or fluctuations in i) content duplication and that in ii) partitioning of biomolecules, particularly in SNAIL (depicted by *f*(*SNAIL^par^*,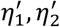)) (**Fig 1C**). (For more information on formalism used to include content duplication and partitioning noise, please refer to Methods section 1). Each cell contains the ZEB1/miR-200 feedback loop driven by SNAIL, and the levels of these molecules define the state of each cell. Depending on the levels of SNAIL, cells may acquire a phenotype among all the stable ones, as shown in the bifurcation diagram (**Fig 1D**) [22]. At SNAIL = 150K molecules, all cells can adopt only an epithelial state (lower blue curve in Fig 1D); at SNAIL = 200K molecules, a cell can acquire any of the three states – epithelial (E; lower blue curve), mesenchymal (M; top blue curve) or hybrid E/M (middle blue curve), while at SNAIL = 250K molecules, all cells adopt a mesenchymal state (top blue curve in Fig 1D). Thus, asymmetry in content duplication and/or partitioning of SNAIL levels can alter the SNAIL values sufficiently enough so as to allow a phenotypic switch. For instance, if one daughter cell has SNAIL = 250K for a parent cell with SNAIL = 200K, then the daughter cell will be mesenchymal in nature irrespective of the phenotype of the parent cell (E, hybrid E/M or M). We have also implemented *in silico* passaging to mimic the experimental protocol for conducting these experiments, where 10% of the cell population is passaged maintaining the distribution of cells in different phenotypes, when the entire population reaches 80% of its carrying capacity (**Fig 1E**). Further, the division rate of each subpopulation of cells is considered to follow logistic growth rate, whereas the death rate is directly proportional to subpopulation size. Together, these factors are incorporated (**Fig 1F**) in a population dynamics model including cell division which may be accompanied with a phenotypic switch (please see Methods sections 2, 3 for more details about population dynamics model).

**Fig 1.**
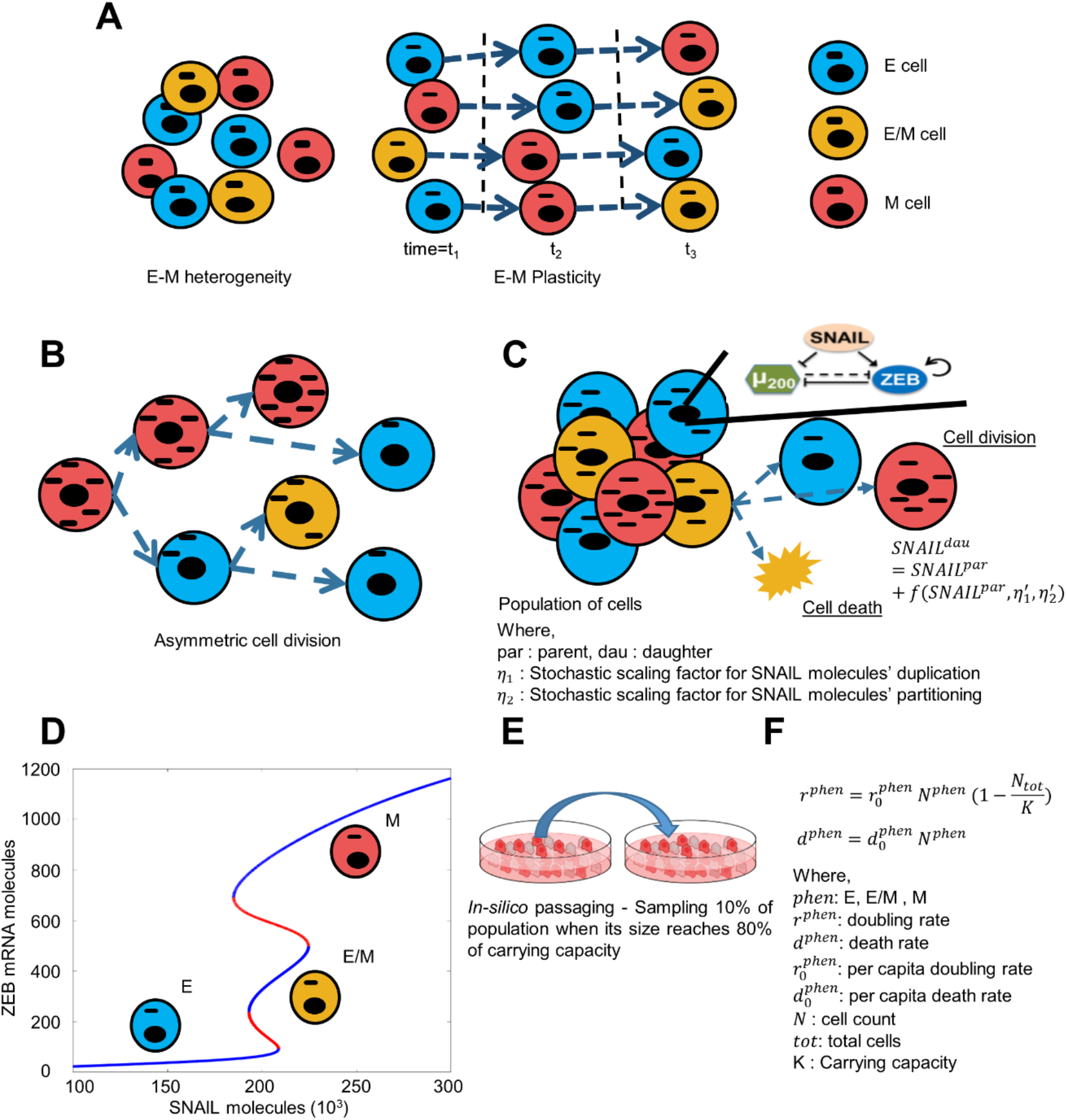
Model description for population growth accompanying E-M heterogeneity. **A)** Schematic showing existing E-M heterogeneity among cancer cells and their spontaneous phenotypic switching. **B)** Schematic for asymmetric distribution of biomolecules among daughter cells as a potential cause of spontaneous phenotypic switching. **C)** Each cell in the population is assigned with a core EMT network. It can divide or die in a given time step depending upon doubling and death rates. When it divides, each daughter cell inherits parent SNAIL levels taking consideration of fluctuations in its content duplication and partitioning during cell division. **D)** Different stable (blue curves) ZEB mRNA levels based on SNAIL levels. Low, medium, and high ZEB mRNA levels corresponds to Epithelial (E), Hybrid (E/M), and Mesenchymal (M) phenotypes respectively. This bifurcation diagram is for the miR-200/ZEB feedback loop driven by SNAIL, as adapted from Lu et al. PNAS 2013 [22]. **E)** Schematic for in-silico passaging; adapted from https://freesvg.org/imacie-of-cell-culture-dish. **F)** Formalism for cell doubling and death rates for all three phenotypes (E, E/M, and M) of cells

Using this framework, we investigated how the population distribution emerged over time as we started with different initial fractions, and whether we can recapitulate the dominance of epithelial (EpCAM^high^) subpopulation over a mesenchymal (EpCAM^low^) one as seen experimentally (**Fig 2A**) [20]. We first experimentally quantified doubling time of PMC42-LA cells to be 22.67 +/− 2.77 hours (**Fig 2A**). We started our simulations with four distinct initial conditions: 1) Epithelial dominated (initial fraction E: E/M: M = 0.7: 0.1: 0.2), 2) Mesenchymal dominated (initial fraction E: E/M: M = 0.2: 0.1: 0.7), Hybrid dominated (initial fraction E: E/M: M = 0.1: 0.7: 0.2), and mixed fractions (initial fraction E: E/M: M = 0.4: 0.2: 0.4). In these simulations, we considered an average doubling time of the population to be 20 hours, *η*_1_ (scaling factor for noise due to SNAIL molecules’ duplication) = 0.2 and *η*_2_ (scaling factor for noise due to SNAIL molecules’ partitioning) = 0.1, and tracked the population distribution as a function of time. This choice of values represents typical noise in protein levels reported over a cell division [24].

**Fig 2.**
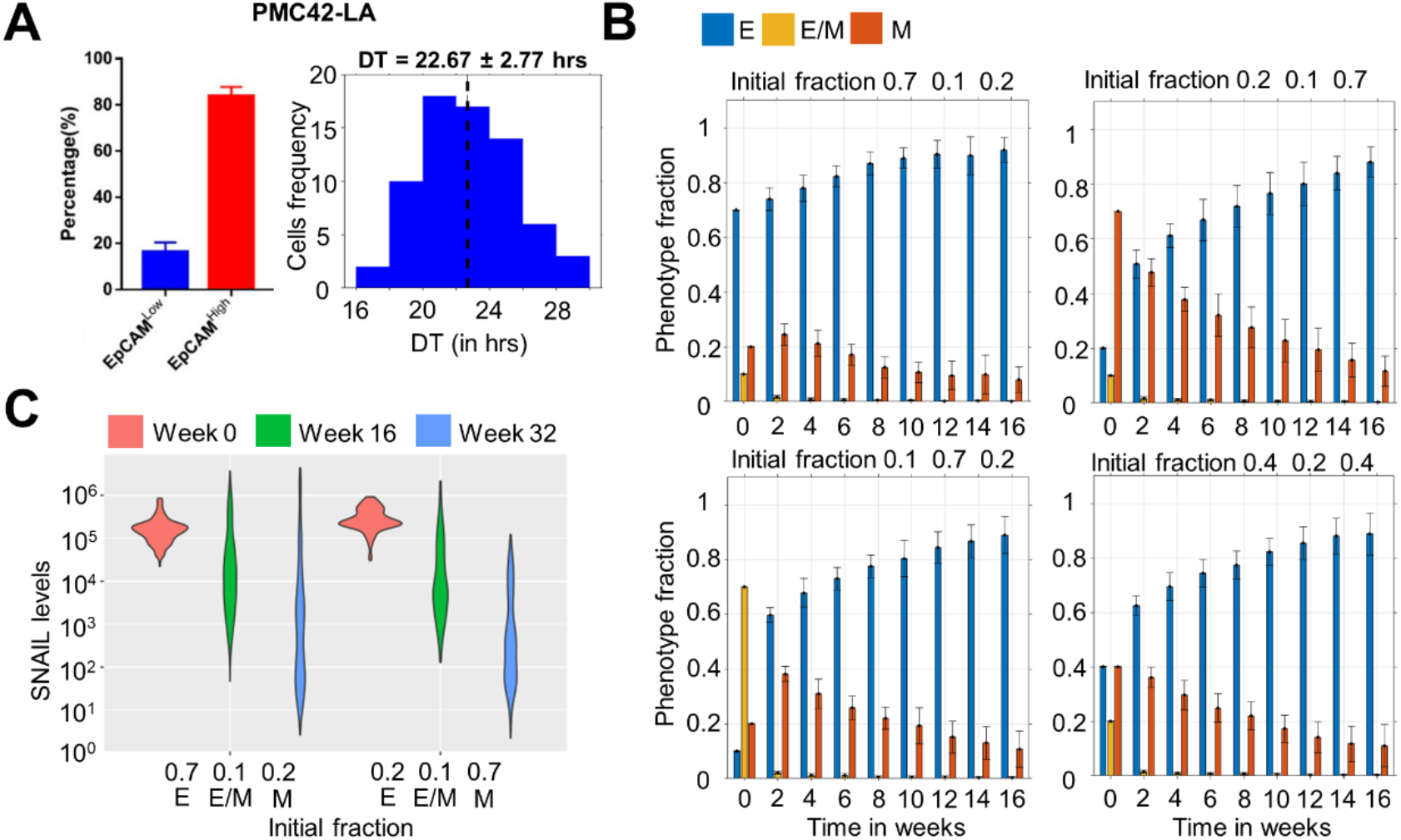
Dominance of epithelial (EpCAM^high^) phenotype in the population over time for multiple initial distributions. **A)** Phenotypic distribution of EpCAM^high^ and EpCAM^low^ subpopulations in PMC42-LA cells (left – adapted from Bhatia et al. [20]), and their observed doubling time distribution (right). **B)** Changes in phenotypic fraction over time starting with different fractions of E, E/M, and M cells in the population. **C)** Change in distribution range of SNAIL levels over 32 weeks for two different initial conditions. Average doubling time (DT) of each phenotype is set to 20 hrs and scaling factors *η*_1_ and *η*_2_ to 0.2 and 0.1. The initial population size was 200 cells. Mean and standard deviation calculated from 16 independent runs.

We observed that all the initial fractions converged to an epithelial dominant population over a period of 16 weeks (**Fig 2B**), i.e. greater than or equal to 80% population being E (EpCAM^high^). Particularly, for mixed initial fraction (E: E/M: M = 0.4: 0.2: 0.4), most of the hybrid E/M cells switch phenotype either to E or M within 2 weeks of time, after which population is mostly comprised of E and M cells, which eventually converges to an epithelial dominant one. Concomitantly, there is also a shift in the distribution of SNAIL levels, such that the range of SNAIL values observed tend to correspond to an epithelial phenotype as well by week 16 and 32, as compared to week 0 (**Fig 2C**), thereby explaining a gain in epithelial-dominated subpopulation as seen experimentally, i.e. EpCAM^high^ subpopulation constituting the majority of PMC42-LA cell line [20]. Further experiments revealed that this population distribution can be reproduced by the FACS-sorted EpCAM^high^ and EpCAM^low^ subpopulations when cultured individually, thus reminiscent of our simulations showing the asymptotic dominance of epithelial subpopulation irrespective of initial phenotypic distributions.

Over longer simulation times in our model, the dominance of epithelial fraction grew even stronger (**Fig S1A**). Thus, while our model encapsulates the dominance of EpCAM^high^ subpopulation in PMC42-LA cells, it cannot accurately reproduce the experimentally observed 80:20 EpCAM^high^: EpCAM^low^ ratio. This lacuna indicates the role of various important factors (both cell-autonomous and non-cell-autonomous: chromatin status and cellular communication, for instance, respectively) which can influence stochastic fluctuations during cell division induced spontaneous switching, thus altering this ratio. Nonetheless, our simple phenomenological model can reproduce salient features of population dynamics reported in the PMC42-LA cell line [20].

To assess how fluctuations in other players of EMT network (miR-200, ZEB) influence population dynamics, we introduced stochasticity in their content duplication and partitioning during cell division, rather than just in SNAIL levels. The population dynamics for this scenario, using the parameters and initial fractions described above, gave qualitatively similar results of epithelial dominance over mesenchymal, though the time taken to gain this dominance was shorter in this case (**Fig S1B-C**), indicating that additional noise can accelerate the system dynamics. These results suggest that accounting for asymmetry in the levels of SNAIL is sufficient to capture the qualitative population dynamics for PMC42-LA cells.

The dominance of epithelial phenotype in a population over time irrespective of initial fractions of phenotypes points towards the possibility that hybrid E/M and mesenchymal cells switch more frequently to epithelial as compared to epithelial states switching to hybrid E/M and mesenchymal. This one-sided higher switching rate can be explained by multiplicative nature of noise considered (lower levels of SNAIL in epithelial cells invoke further less fluctuations during division) and have been quantified later (**Fig 3**).

**Fig 3.**
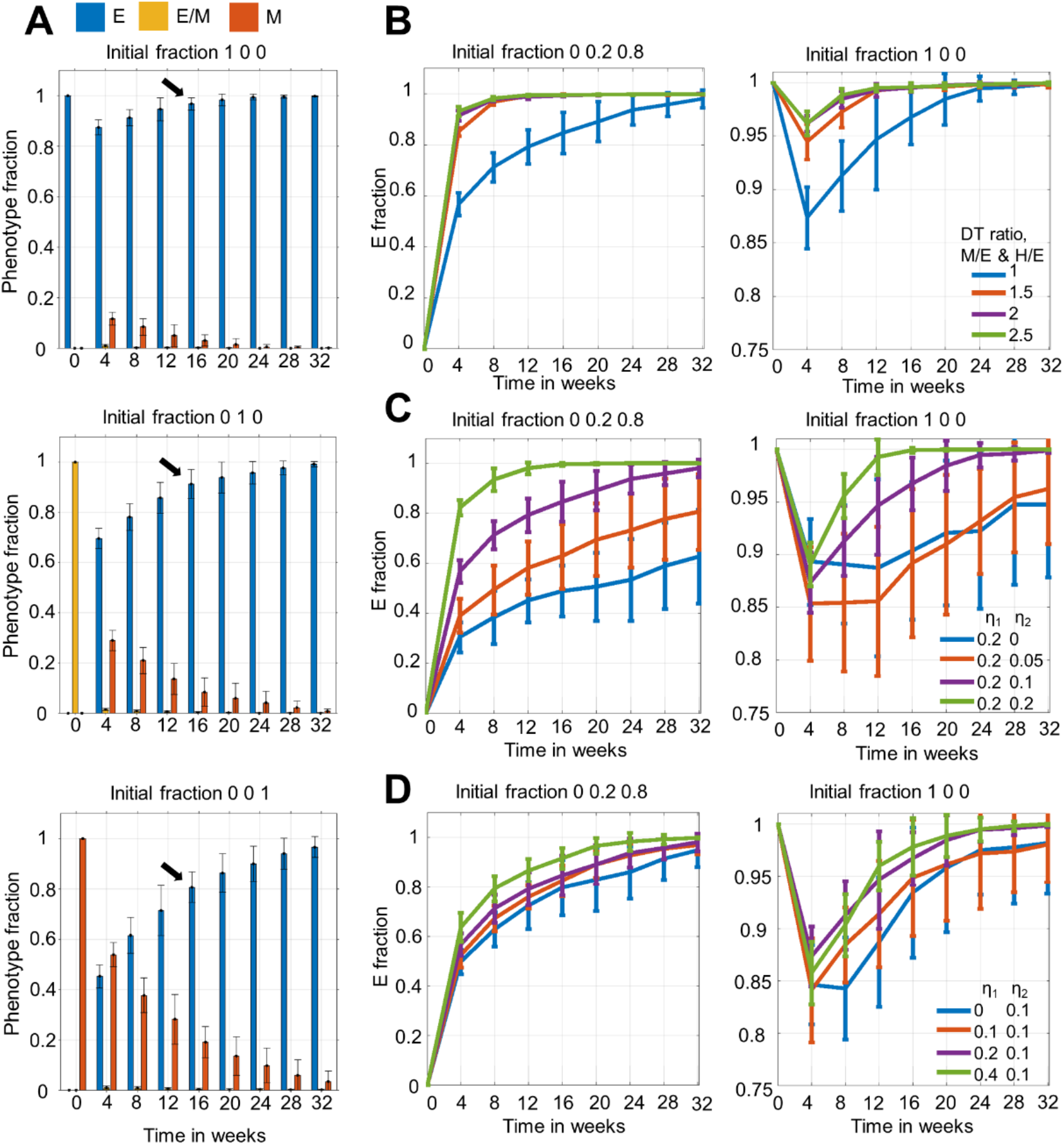
Time to attain dominance of E cells depends on initial fraction and average doubling times of phenotypes, and the extent of molecular fluctuations. **A)** Temporal changes in phenotypic distribution for purely E, E/M, and M initial population. **B)** Temporal changes in phenotypic distribution when there is heterogeneity in avg. doubling time (DT) among phenotypes. DT(E/M, M) = DT ratio * DT(E). *η*_1_ = 0.2, and *η*_2_ = 0.1. **C)** Temporal changes in phenotypic distribution for fixed *η*_1_ and varying *η*_2_ values. Here, DT(E, E/M, M) = 20 hrs. **D)** Same as Fig3C but with varying *η*_1_ and fixed *η*_2_ values. In all, except (A), initial fractions of 1) Mix of E/M and M, and 2) pure E phenotypes are considered. The initial population size was 200 cells. Mean and standard deviation calculated from 16 independent runs.

### Time to attain epithelial dominance depends on initial fractions, doubling times of phenotypes and the extent of stochastic fluctuations during cell division

Upon simulating the population dynamics starting with purely E and purely M phenotypes, we noticed differences in mean epithelial (E) fraction at week 16 (**Fig 3A**). When starting with purely M phenotype, it took 16 weeks to arrive at an epithelial-dominant population as compared to starting with hybrid E/M (8-12 weeks) or fully E ones. This trend raised the possibility that while initial phenotypic distribution may not alter the steady state itself, it can change the time taken to arrive at it. For the scenario starting with purely hybrid E/M cells, within 4 weeks the population structure had approximately 60-70% epithelial cells, thus its dynamics post the 4 week timepoint is understandably similar to that seen for E: E/M: M = 0.7: 0.1:0.2 scenario (compare middle panel in **Fig 3A** with the top left panel in **Fig 2B**).

Besides initial phenotypic distribution, another factor that can impact the population dynamics is average doubling time (DT). So far, we considered the same DT for E, E/M, and M phenotypes. But experimental evidence suggests slowing down of proliferation rate of cells on undergoing EMT [20,25]. Thus, we considered the case of increased DT during EMT, by keeping the average DT of E/M and M phenotypes as 1.5, 2, and 2.5 times more than that of E phenotype. We observed that the population maintained its epithelial dominance, and converged to a stable phenotypic distribution faster than the case of when all cells doubled at the same rate (**Fig 3B**), irrespective of the initial condition. This trend can be explained by a higher resilience of E cells to switch to a hybrid E/M or M phenotype during cell division, now coupled with their higher proliferation rate, thereby offering the epithelial subpopulation an additional advantage to amplify their population fraction. Importantly, this trend was already seen at the DT of hybrid E/M and M cells being 1.5 times than that of the E cells, hence indicating that a 50% increase in doubling time for cells undergoing EMT may be sufficient in influencing the population structure. The slight initial decrease in epithelial fraction noticed for the purely E case (**Fig 3A, B**) can be explained by appreciating that SNAIL levels for the initial cell population were sampled from log-normal distribution, whose median was centred on the SNAIL level where all phenotypes were stable (tristable region in Fig 1D) and therefore, the cells were highly susceptible to undergo phenotypic switching within the first few cell divisions. Despite this initial dip, an epithelial dominant population emerged eventually.

Next, we investigated how the extent of stochastic fluctuations in SNAIL molecules being doubled and partitioned (*η*_1_ and *η*_2_ respectively) influenced phenotypic distribution over time. When we varied *η*_2_, while maintaining the values of *η*_1_ = 0.2 and average population DT = 20 hours, we noticed that for the mesenchymal dominated initial fraction (E: E/M: M = 0: 0.2: 0.8), the fraction of epithelial cells was higher for a higher *η*_2_ value for the same time point (**Fig 3C**, left). However, not much observable effect on this fraction was noticed when starting with an epithelial dominated population (**Fig 3C**, right). Similar observations were made when we varied *η*_1_ instead of varying *η*_2_ (**Fig 3D**). Thus, amplifying fluctuations in either duplication or partitioning of SNAIL molecules seemed to enhance the chance of phenotypic switching for a M cell much more than for an E cell.

When we accounted for heterogeneity in average DT along with increasing fluctuations in SNAIL levels during cell division, the fast proliferating and relatively stable E cells grew much faster than the slow proliferating and more plastic E/M and M cells, enriching for epithelial cells (**Fig S2A-D**).

Finally, we analysed how the population dynamics was altered when average DT was increased for all phenotypes, given the experimentally observed average DT can often depend on confluency of cells in a petri dish. Thus, for this, we simulated population dynamics keeping average DT of all three phenotypes as 30 hrs, which led to overall slower dynamics (**Fig S2E**). Instead of plotting against absolute time units, we also took number of cell cycles as the x-axis, whose one unit is the population’s average DT. This helped to compare the overall changes in phenotypic fractions between DT of 20 and 30 hrs scenarios. We found that given an equal number of cell cycles, the changes in E fractions were similar (**Fig S2F**). These observations help us to conclude that even if all cells, on an average, divided slower, the population growth and phenotypic switching trajectory would be similar to when cells divided faster, when normalized with average DT.

### Phenotypic switching probability and rate in cell division events depends on the cells’ location on E-M axis

After characterizing the population dynamics at various time points as a function of different model parameters, we wanted to better understand it from a cell division perspective. In our framework, a cell can undergo one of the three division types: 1) symmetric division – when both daughter cells have same phenotype as the parent cell, 2) asymmetric division – when one daughter has phenotype different than parent, and 3) divergent division – when both daughters have phenotype different than parent (**Fig 4A**). To quantify the probability of cells undergoing one of the three division types, we analysed certain cells occupying possible stable phenotypes spread across the SNAIL ranges (**Fig 4B**). Iterating cell division events at a given SNAIL value, we tracked the phenotypes of daughter cells, at specific *η*_1_ and *η*_2_ values, thus calculating different division probabilities over an ensemble of iterations. At *η*_1_ = 0.2, *η*_2_ = 0.1, in SNAIL levels regions where either E or M phenotypes were the only stable state (monostable regions in bifurcation diagram – **Fig 4B**; SNAIL = 100K, 300K), more than 90% events were of symmetric division. However, as the SNAIL levels corresponded to multi-stable region (SNAIL values =150K, 250K), there was an increasing tendency to undergo asymmetric division, which was higher for M than for E cells. In different bi-stable regions (SNAIL = 189K for {E, M} and SNAIL = 219K for {E/M, M}), with further increasing probability of asymmetric division, divergent division also became more prominent and was the most dominant division for hybrid E/M cells (**Fig 4C**). This trend explains the sudden drop in hybrid E/M fraction of population to very low levels within two weeks, when starting from a hybrid dominant population (**Fig 3A**). Further, in the tri-stable region (SNAIL = 206K for {E, E/M, M}), the probabilities for both divergent and asymmetric division were increased (**Fig 4C**). Put together, the probability of phenotypic switching at cell division is the highest in tri-stable region (intermediate SNAIL levels ~200K) and decreases for cells as their corresponding SNAIL levels either increase or decrease.

**Fig 4.**
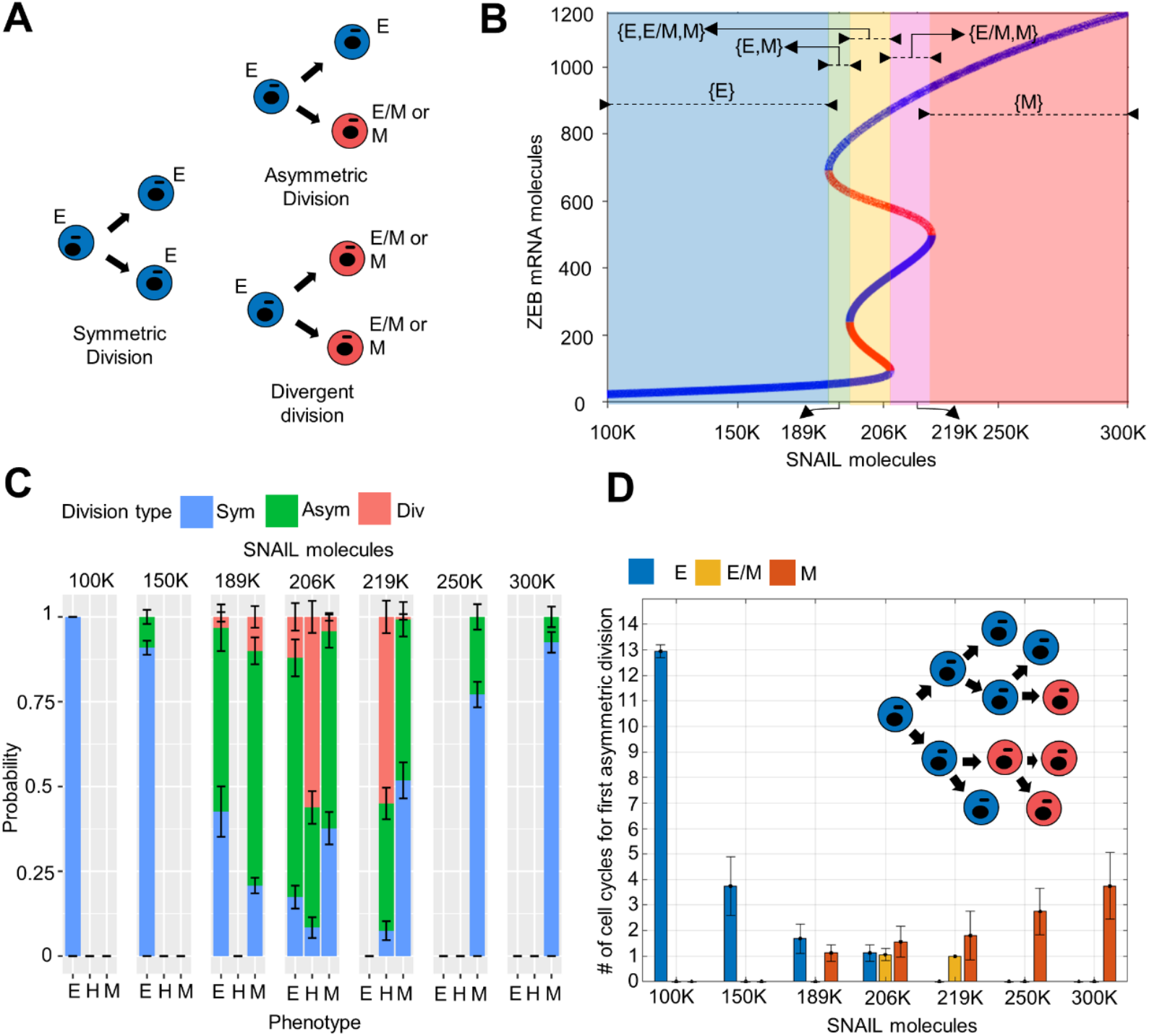
Phenotypic switching probability and its rate at cell division depends on the location of a cell on E-M axis. **A)** Schematic of different possible cell division types. **B)** Different ranges of SNAIL where E, E/M, and M phenotypes are stable. **C)** Probabilities of an E, E/M or M cell to undergo one among the three division types (Fig 4A) when its SNAIL levels lie in different regions in bifurcation diagram (Fig 4B). **D)** Number of cell cycles (generations) required to make first asymmetric or divergent division when a parent E, E/M or M cells’ SNAIL level lied in different regions in bifurcation diagram (Fig 4B) (schematic given in inset). Mean and standard deviation were calculated from 10 and 16 independent runs in (C) and (D), respectively. In C), each run includes 100 iterations. *η*_1_ = 0.2, *η*_2_ = 0.1.

Next, we quantified these probabilities for varying *η*_1_ and *η*_2_ values. While *η*_1_ = 0 resulted in either asymmetric or symmetric division of E and M cells across SNAIL levels (i.e. preventing divergent division), *η*_2_ = 0 leads to only symmetric and divergent divisions for these two phenotypes (i.e. preventing asymmetric division) (**Fig S3A,B**). Also, higher values of *η*_1_ and *η*_2_ amplify the chances of divergent and asymmetric division, respectively, across SNAIL ranges (increasing *η*_1_ in **Fig S3A,C,E** and increasing *η*_2_ in **Fig S3B,D,F**). Thus, *η*_1_ and *η*_2_ - the factors that represent noise during cell division - can alter the probabilities of undergoing symmetric, asymmetric and divergent division types for a cell with a SNAIL level (**Fig S3**).

We observed that cells with SNAIL levels well away from the multi-stable phenotypic regions have mostly undergone symmetric division. However, when we started with such a homogenous or largely homogeneous population and tracked the phenotypes of daughter cells over multiple cell cycles, we noted phenotypic switching in which at least one daughter cell took a different phenotype (**Fig 3**). Thus, we quantified how many cell divisions it took for a cell to give rise to one of the cells in its progeny with a different phenotype than its own. We observed the progeny up to 12 generations/cell cycles. We saw that the cells with SNAIL levels in a multi-stable region switch phenotype within one or two cell cycles (**Fig 4D**). We also noticed a skew between the resilience of E and M cells to phenotypic switching in their mono-stable regions, i.e. E cells required more cell cycles to give rise to a non-similar progeny cell than the M cells did (compare the behavior seen at S=300K and S=250K with that at S=100K and S=150K in **Fig 4D**). This difference may underlie the phenomenon of E cells dominating over E/M and M cells in the population over time. However, this skew vanished in bi-stable and tri-stable regions, where all three phenotypes became equally susceptible to undergo asymmetric switching within few generations/cell cycles (S=189K, 206K, 219K in **Fig 4D**).

We also examined how *η*_1_ and *η*_2_ values varied the number of cell cycles over which progeny diversification was observed. Increase in either *η*_1_ and *η*_2_ caused faster switching for all phenotypes of cells, i.e. less number of cell divisions was required, on average, for a cell to give rise to a different phenotype (increasing *η*_1_ in **Fig S4A-C** and increasing *η*_2_ in **Fig S4D-F**). However, *η*_2_ contributed more as compared to *η*_1_ (compare **Fig S4A-C** with **Fig S4D-F;** quantified in **Fig S5,S6**).

### Heterogeneity in E fraction at initial time points among single cell clones

So far, we have focused on population dynamics when starting with an initial cell population; however, heterogeneity has also been observed experimentally in single-cell clones established from cell lines [20,21]. For instance, in distinct single-cell clones established from PMC42-LA, varying distributions of EpCAM^high^: EpCAM^low^ subpopulations were seen after initial two passages (**Fig 5A**) [20]. We interrogated whether our model can reproduce this heterogeneous behavior.

**Fig 5.**
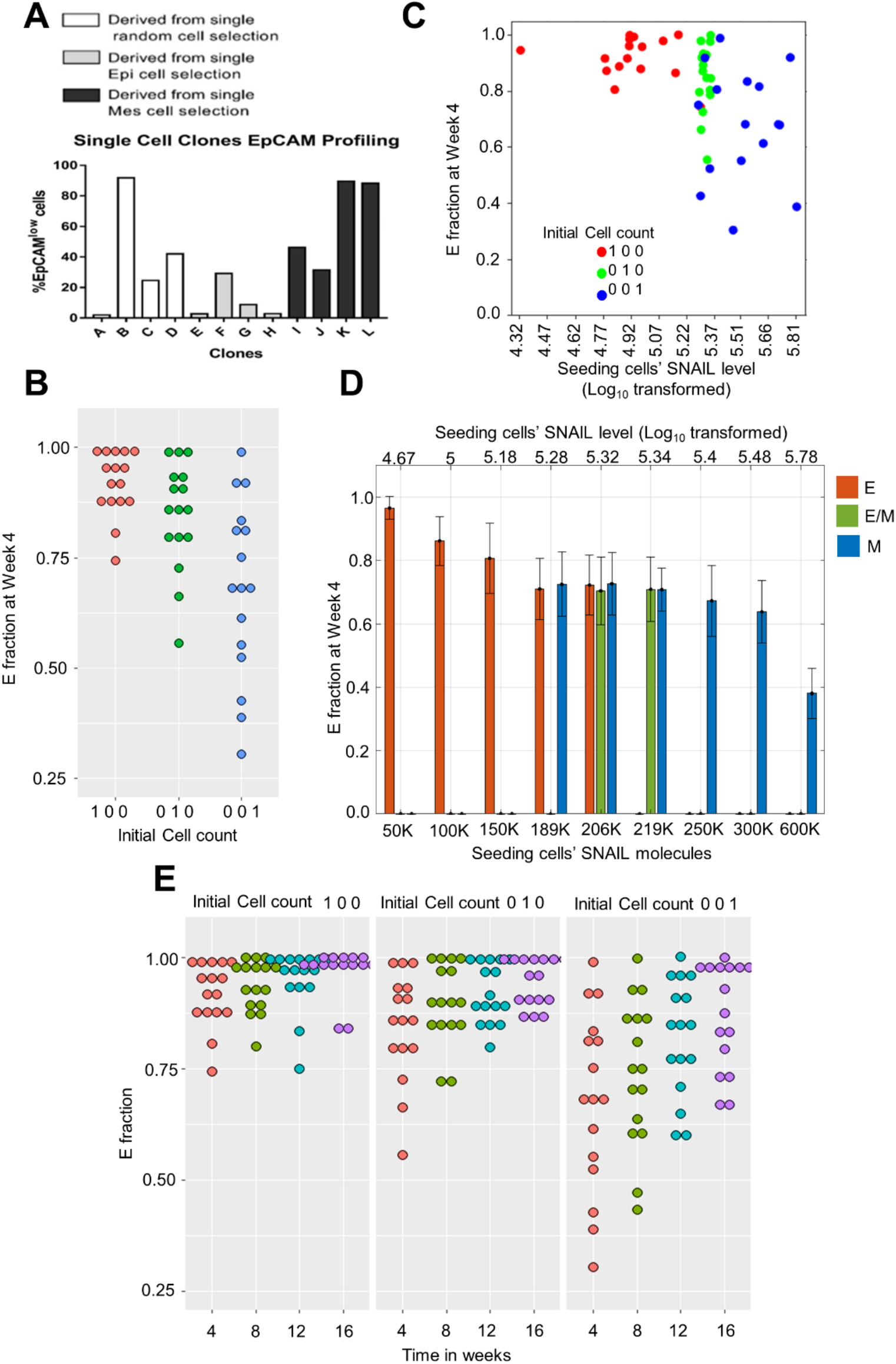
Heterogeneity in E fraction among single cell clones at initial stages of culture. (A) Experimentally observed heterogeneity in EpCAM profiling of single cell clones from PMC42-LA cells (adapted from Bhatia et al. [20]). (B) Variability in E fraction observed on simulating population dynamics starting with single E, E/M and M cell. Each dot represents E fraction at 4^th^ week in an independent single cell simulation run. (C) E fraction in a single cell clone at 4^th^ week plotted against the seeding (parent) cells’ SNAIL level. (D) Spread of E fraction at 4^th^ week of single cells clones when the seeding cells were initialised with certain SNAIL levels spanning mono-, bi-, and tri-stable regions of bifurcation diagram (Fig 4B). Mean and standard deviation calculated from 20 independent runs (E) Temporal dynamics extension of single cell simulation runs in Fig 5B. All results are with *η*_1_ = 0.2 and *η*_2_ = 0.1, and DT(E, E/M, M) = 20 hrs.

We performed population dynamics simulation starting will single E, E/M and M cell, maintaining *η*_1_ = 0.2, *η*_2_ = 0.1, and average DT as 20 hours. We observed heterogeneity in E fraction at week 4 when multiple such single cell simulation runs were performed (**Fig 5B**). Thus, as a proof of principle, our model could recapitulate the experimentally observed heterogeneity in EpCAM^low^ fraction. In these simulations for single-cell clones, at week 4 time point, the heterogeneity in fraction of E cells was the highest when the seeding cell was mesenchymal (M). Among the M clones, the highest E fraction noticed was close to the highest E fraction noticed for single-cell clones established from E or E/M initial phenotype. However, in M clones, we observed instances where the E fraction was as low as 27% (**Fig 5B**).

To examine this heterogeneous behaviour of individual clones more closely, we probed the levels of SNAIL in the seeding (individual) cell for each of these clones. This led us to identify the range of SNAIL levels in the individual cells that were all *‘cultured’ in silico* to develop a clone (**Fig 5C**). From this range, we identified representative SNAIL values and independently established singlecell clones from them. Interestingly, the single-cell clones showed heterogeneity in E fraction at 4 weeks, despite being seeded with the same SNAIL level (**Fig 5D**), reminiscent of stochastic effects at lower (cell) numbers. Also, as expected, the average E fraction decreases as seeding SNAIL levels are increased (compare the average E fraction at SNAIL= 600K vs. that at SNAIL= 50K in Fig 5D). Another feature we noticed is that the clones established from cells with the same initial phenotype (M) but different initial SNAIL levels had varying E fractions at week 4 (compare the average E fraction at SNAIL = 206K vs. that at SNAIL = 600K in Fig 5D). This extent of diversity is less when the initial cell belongs to an epithelial or a hybrid E/M phenotype. This difference between the extent of variability noticed can possibly explain why we see more heterogeneity in E fraction when starting from initially mesenchymal cell as compared to an initially epithelial or hybrid E/M one (**Fig 5B-D**).

Finally, when we continued the single-cell (clonal) simulations for longer duration, we observed a decrease in heterogeneity in the E fractions with time (**Fig 5E**). Further, the E fraction for all single cell clones increased overall. This difference in short-term vs. long-term behavior can be possibly rationalized by our population dynamics simulations earlier showing predominance of epithelial phenotypes irrespective of initial phenotypic distributions (**Fig 2B, 3A**), if we consider the clonal distribution noticed at week 4 as the initial condition for simulations being continued until week 16 or later.

## Discussion

Understanding the molecular mechanisms underlying epithelial-mesenchymal plasticity and heterogeneity can contribute to better therapeutic strategies [26]. These mechanisms can be context-specific with varying degrees of contribution to genetic and/or non-genetic heterogeneity. Epigenetic alterations, for instance, can govern the rate of bidirectional switching among the phenotypes, enabling reversible or irreversible EMT, as well as driving resistance to undergo EMT [27–29]. Cell-cell communication through autocrine and/or paracrine signaling with other tumor cells as well as stromal cells can also shape the E-M phenotypic heterogeneity patterns in a population [19,30–32]. Another contributing factor can be differential activation of many signaling pathways implicated in EMT, thus generating a varied phenotypic repertoire of states in the multidimensional EMT landscape [33–37]. Here, we highlight one other possible reason driving E-M heterogeneity – phenotypic switching due to asymmetric cell division driven by noise in the processes of content duplication and in partitioning of biomolecules. We investigate the influence of such fluctuations on levels of SNAIL – a driver of miR-200/ZEB feedback loop – during cell division in determining the phenotypic distribution of population, but our framework is applicable to investigate the population dynamics emerging from stochastic partitioning of molecules involved in other multi-stable EMT networks [38,39] as well.

Asymmetric cell division is an evolutionarily conserved mechanism used by prokaryotes as well as eukaryotes to generate cell-to-cell heterogeneity, and mediate cell-fate decisions [40,41]. Not only biomolecules (RNAs, proteins), but also entire organelles such as mitochondria and endoplasmic reticulum can be asymmetrically partitioned, with implications in cancer cell proliferation rates [42]. This phenomenon has been observed in multiple cancers [43–45], but its functional consequences remain largely unexplored. Although our modeling framework does not yet specifically incorporate molecular mechanisms regulating this phenomenon [40], our results suggest that one possible consequence of fluctuations during cell division can be phenotypic switching and heterogeneity among subpopulations. Recent reports in glioblastoma have demonstrated that asymmetric enrichment of EGFR and p75NTR in a daughter cell during cell division conferred enhanced resistance to the standard-of-care therapies such as radiation and temozolomide [46]. While we do not yet know about differences, if any, in the drug resistance features of EpCAM^high^ and EpCAM^low^ sub-populations in PMC42-LA cells, the varied drug-resistance features seen in singlecell clones established from PMC42-LA [20] can be a putative outcome of underlying asymmetric cell division. Approximately 10-30% of cells undergoing TGFβ-driven EMT were seen to exhibit asymmetric cell division, as traced by NUMB distribution in daughter cells [47], but whether this asymmetry led to phenotypic switching was not tracked *per se*. Therefore, our model suggests that blocking cell division can be a possible way to restrict phenotypic plasticity and/or heterogeneity. Preliminary experimental observations made recently support this prediction by our model [47].

Our model can recapitulate the observations for PMC42-LA system, an epithelial-dominant subline. However, what mechanisms may explain phenotypic heterogeneity in a mesenchymal-dominant population, such as EM3 or M clone from SUM149 cell line [21], remains to be investigated further. One factor that can alter the model outcomes is the way noise during cell division is incorporated. We have considered multiplicative noise (fluctuations in SNAIL proportional to its levels); however, our previous effort encapsulating additive noise (constant magnitude of fluctuations in SNAIL, irrespective of its levels) could explain spontaneous phenotypic switching observations in prostate cancer *PKV* cell line [48]. Whether cancer cells exhibit additive or multiplicative noise during cell division remains unknown experimentally. Further, this noise and/or its consequences can be influenced by mutually dependent factors, such as chromatin status and diffusible cytokines [49]. These factors have not yet been explicitly incorporated in our framework.

Despite the above-mentioned limitations, our model recapitulates various observations for the PMC42-LA system: a) stable dominance of the EpCAM^high^ subpopulation, b) repopulation of parental distributions starting with only one subpopulation, and c) enhanced heterogeneity in EpCAM^high^: EpCAM^low^ ratio of cells in single-cell derived clones. We predict that these single-cell derived clones converge to EpCAM^high^ dominant distribution in longer time-scales, a prediction which remains to be experimentally verified. Thus, we demonstrate that during cell division, stochasticity in content duplication and partitioning of molecules involved in EMT can lead to spontaneous state switching, and therefore generate non-genetic heterogeneity.

Future efforts are directed towards integrating continuous stochastic fluctuations in EMT drivers with asymmetric cell division which happens at discrete time-steps [50]. Addressing these questions will involve mathematical models that can decode the emergent dynamics at multiple scales – regulatory levels (transcriptional, epigenetic), length (intracellular, non-cell-autonomous effects by cytokines) and time (cell division, chromatin remodelling, stochastic gene expression).

## Author contributions

PJ performed simulations, SB performed experiments, EWT and MKJ conceived and supervised research. All authors contributed to data analysis and writing of the manuscript.

## Code Availability

All codes used are available at https://github.com/Paras-Jain20/EMT-Population-Dynamics

## Methods

### 1. Asymmetric distribution of molecular content on cell division

Following the method proposed earlier [48], we consider fluctuations in the levels of cellular content during its inheritance by daughter cells on cell division. These fluctuations arise due to both imperfect duplication during cell cycle and later asymmetric partitioning to the daughter cells. We propose these fluctuations to be proportional to the amount of the molecular content available in the dividing parent cell itself. Considering 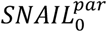 denotes SNAIL level in a cell right after its division. Now, during cell cycle SNAIL content will approximately get doubled, so that right before next cell division we can write:

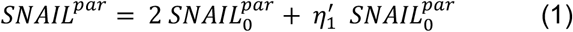

Where, 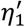 is a stochastic scaling factor that determine the fluctuation due to imperfect molecule duplication.

Next, when a parent cell partitions its molecular content to two daughter cells during cell division, SNAIL levels in each daughter can be specified as:

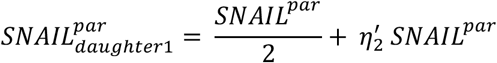

On substituting *SNAIL^par^* from (1),

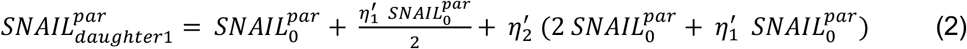

And, for the other daughter cell

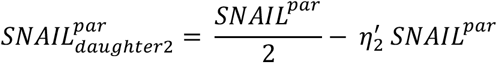

On substituting *SNAIL^par^* from (1),

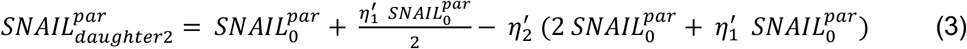

Where, 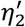 another random scaling factor determining the fluctuation in SNAIL levels due to partitioning at the time of cell division.

We consider stochastic scaling factors 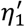 and 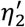 to be two independent normally distributed random variables with zero means and *η_1_* and *η_2_* as standard deviations, i.e.,

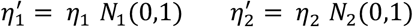

Where, *N_i_*(0,1), *i* = 1,2 represents standard normal random variable. Hereafter, *η_1_* and *η_2_* are reffered to as scaling factors for noise in SNAIL molecules’ duplications & partitioning, respectively. Thus, equations (2) and (3) can be rewritten as

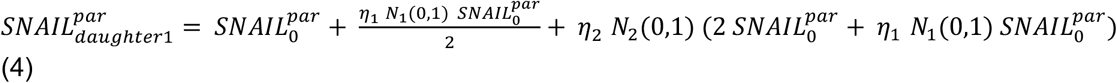

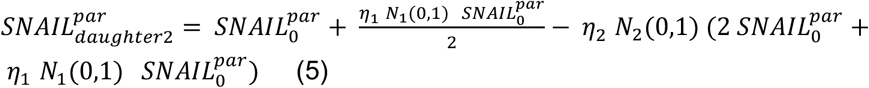

Equations (4) and (5) are used to assign SNAIL values to the daughter cells when a cell division happens. Further, same equations were used when stochastics effects were included in the other players of EMT network (ZEB, mZEB and miR200) at the time of cell division.

### 2. Dynamics of core EMT regulatory network

The dynamics of a core regulatory circuit involving interaction in canonical epithelial (miR200) and mesenchymal (mRNA ZEB and ZEB protein) markers was modelled to explain EMT and MET, based on SNAIL levels [22]. miR200 and ZEB (mRNA and protein) mutually repress each other, and SNAIL supress miR200 levels and activates ZEB at mRNA level. The steady state response of the circuit were analysed for a relevant biological parameter set, which gave bifurcation diagram showing distinct possible stable ranges of ZEB and miR200 based on SNAIL levels as shown in Fig1D. The ordinary differential equations describing the regulation dynamics and model parameters have been described in [22]. The system’s ODEs are listed below:

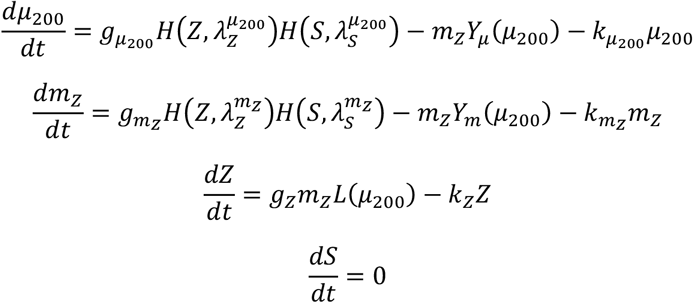

Here, *μ_200_* = [miR-200], *m_z_* = [ZEB1 mRNA], *Z* = [ZEB1], and *S* = [SNAI1]. [·] represents the concentration of a molecular species within a cell. *H* is the shifted Hill function.

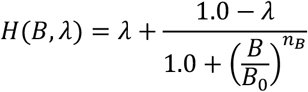

The functions *Y_μ_*, *Y_m_*, and *L* describe the post-transcriptional regulation of mRNA activity by micro-RNAs and have been described in [22].

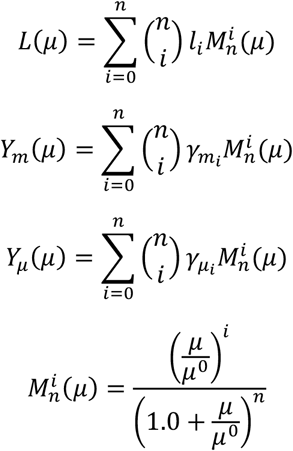

Here, *μ* is the concentration of the micro-RNA and *n* is the number of micro-RNA binding sites on the mRNA. For the inhibition of *ZEB1* mRNA by miR-200, *n* = 6 and 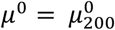. The values of all kinetic parameters are listed in tables S1 and S2.

### 3. Simulation of population dynamics

#### i. Generation of population as per initial phenotypic fraction

Each cell in the system is represented by a set of four variables which hold the levels of miR200, mRNA ZEB, ZEB protein and SNAIL protein for that cell. Random SNAIL values are sampled from a log-normal distribution with median 200×10^3^ and coefficient of variance 1 and all possible stable states corresponding to that SNAIL value are used to initialize the cells’ variables. For example, if sampled SNAIL = 200K and as for this value all three phenotypes – E, E/M and M – are stable. So, three cells are initialized with steady state values of all variables corresponding to each phenotype. Initialization of a cell from a phenotypic state is stopped when its required count in the population is achieved.

#### ii. Avg. birth and death rate of cells

In the cell population, the division rate of cells of a particular phenotype follows the logistic equation shown below:

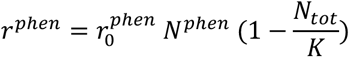

And the death rate of cells of a particular phenotype is as follows:

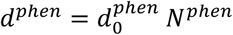

Where,

*phen*: E, E/M, M

*r^phen^*: avg. doubling rate

*d^phen^*: death rate

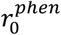: max. avg. doubling rate of an individual cell

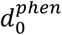: avg. death rate of an individual cell

*N^phen^*: total cells of a phenotype

*N^tot^*: total cells in the population

K: Carrying capacity of the system

#### iii. Population growth

The population growth is simulated using Gillespie’s Stochastic Simulation (SSA) algorithm [51], where six events – three division and three death events for each phenotype are considered. The propensity of occurrence of an event is determined by its average rate as described above. The SSA algorithm tells what next event will be and at what time point. Now, if next event is division of a cell of E phenotype and will occur at t_1_ time, then a cell is uniformly sampled from the pool of cells of that phenotype, and its molecular levels are updated using ODEs for the time gap (t_1_-t_0_), where t_0_ is time point of last most recent event. Then, a new cell is initialized in the population with molecular levels same as that of parent E cell, but with perturbed SNAIL levels. Similarly, the parent cell SNAIL levels are perturbed to account for second daughter cell on division. For a cell death of a phenotype, a cell is uniformly sampled from the pool of cells of that phenotype, and it is erased from the population. Molecular levels of all the other undivided/unaffected cells are updated using ODEs for the time gap (t_1_-t_0_).

### 4. Cell doubling quantification

Images for PMC42-LA cells were captured on PhaseFocus LiveCyte Image Scanner (**Phase Focus**, Sheffield, UK) with 10x magnification; individual images were captured every 11 minutes for a span of 48 hours. Imaging selected regions of interest (ROI) were 750×750μm. 60 individual selected cells were randomly selected and then manually tracked from cytokinesis of a cell to two daughter cells to next cytokinesis to determine the exact cell doubling time.

**Table S1.**

| *Parameter* | *Value* | *Parameter* | *Value* |
| --- | --- | --- | --- |
| $g_{\mu_{200}}$ | $2.1 \times 10^3 \text{ mol. h}^{-1}$ | $n_z^{\mu_{200}}$ | 3 |
| $g_{m_z}$ | $11.0 \text{ mol. h}^{-1}$ | $n_s^{\mu_{200}}$ | 2 |
| $g_z$ | $0.1 \times 10^3 \text{ h}^{-1}$ | $n_z^{m_z}$ | 2 |
| $k_{\mu_{200}}$ | $0.05 \text{ h}^{-1}$ | $n_s^{m_z}$ | 2 |
| $k_{m_z}$ | $0.5 \text{ h}^{-1}$ | $\lambda_z^{\mu_{200}}$ | 0.1 |
| $k_z$ | $0.1 \text{ h}^{-1}$ | $\lambda_s^{\mu_{200}}$ | 0.1 |
| $Z_0^{\mu_{200}}$ | $220.0 \times 10^3 \text{ mol.}$ | $\lambda_z^{m_z}$ | 7.5 |
| $S_0^{\mu_{200}}$ | $180.0 \times 10^3 \text{ mol.}$ | $\lambda_s^{m_z}$ | 10.0 |
| $Z_0^{m_z}$ | $25.0 \times 10^3 \text{ mol.}$ | $\mu_{200}^0$ | 10000 |
| $S_0^{m_z}$ | $180.0 \times 10^3 \text{ mol.}$ | | |

**Table S2.**

| *No. of miRNA binding sites* | *0* | *1* | *2* | *3* | *4* | *5* | *6* |
| --- | --- | --- | --- | --- | --- | --- | --- |
| $l_i$ | 1.0 | 0.6 | 0.3 | 0.1 | 0.05 | 0.05 | 0.05 |
| $\gamma_{m_i} (\text{h}^{-1})$ | 0.0 | 0.04 | 0.2 | 1.0 | 1.0 | 1.0 | 1.0 |
| $\gamma_{\mu_i} (\text{h}^{-1})$ | 0.0 | 0.005 | 0.05 | 0.5 | 0.5 | 0.5 | 0.5 |

Here, $\text{mol.} \equiv \text{molecules / cell}$

## Supplementary figures

**Fig S1.**
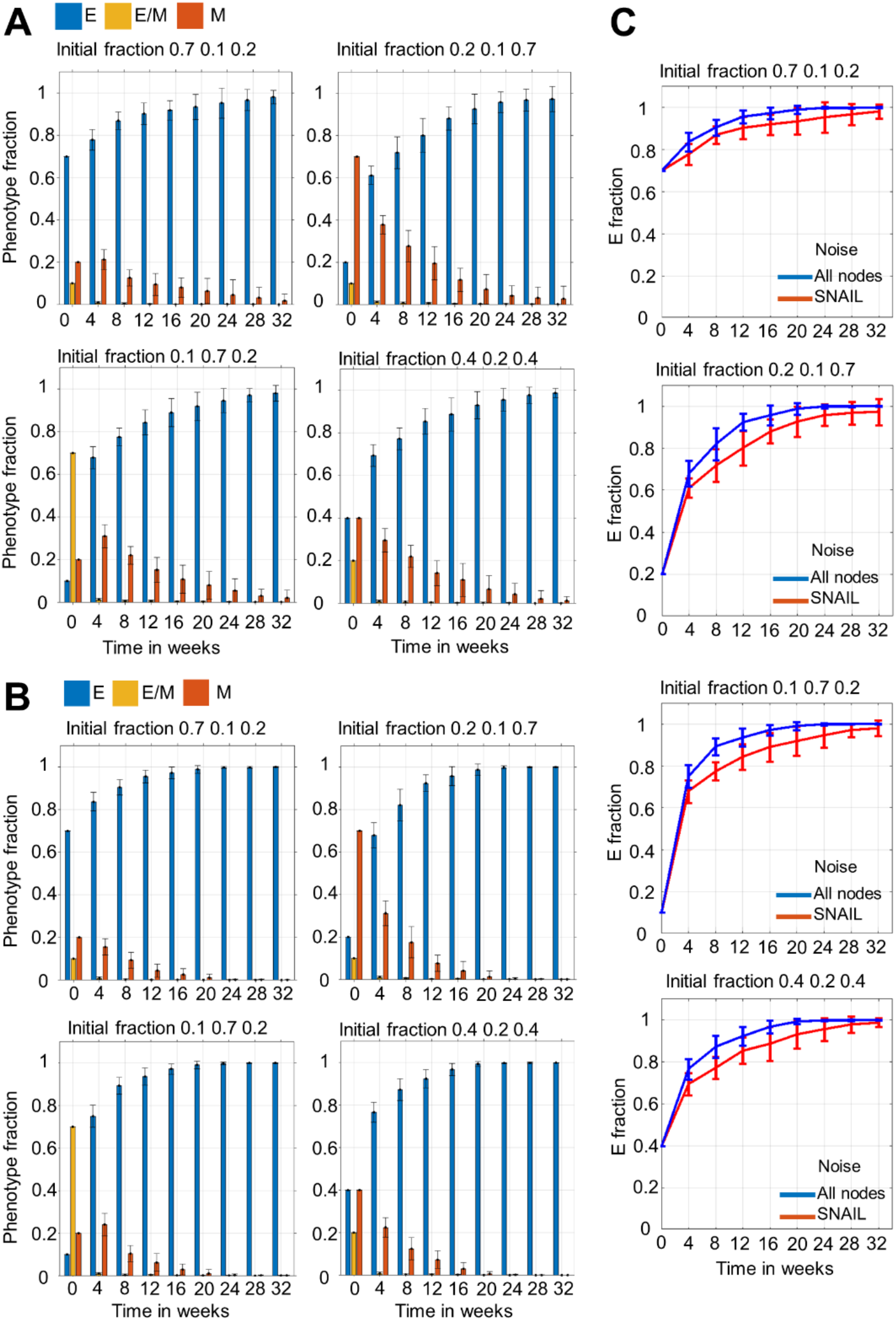
Longer time simulations and similarities between effects of fluctuation in SNAIL and all players in core EMT network during cell division. (A) Extension of simulation of results shown in Fig 1 C to 32 weeks. (B) Temporal changes in phenotypic distribution for same initial fractions as in Fig 1A but with stochastic fluctuations considered in all players of core EMT network during cell division. (C) Overlap in temporal changes in E fraction when SNAIL (red), and all players of core EMT network (blue) had fluctuations in their levels during cell division. Avg. doubling time (DT) of each phenotype is set to 20 hrs and scaling factors *η*_1_ and *η*_2_ to 0.2 and 0.1, respectively. The initial population size was 200 cells. Mean and standard deviation calculated from 16 independent runs.

**Fig S2.**
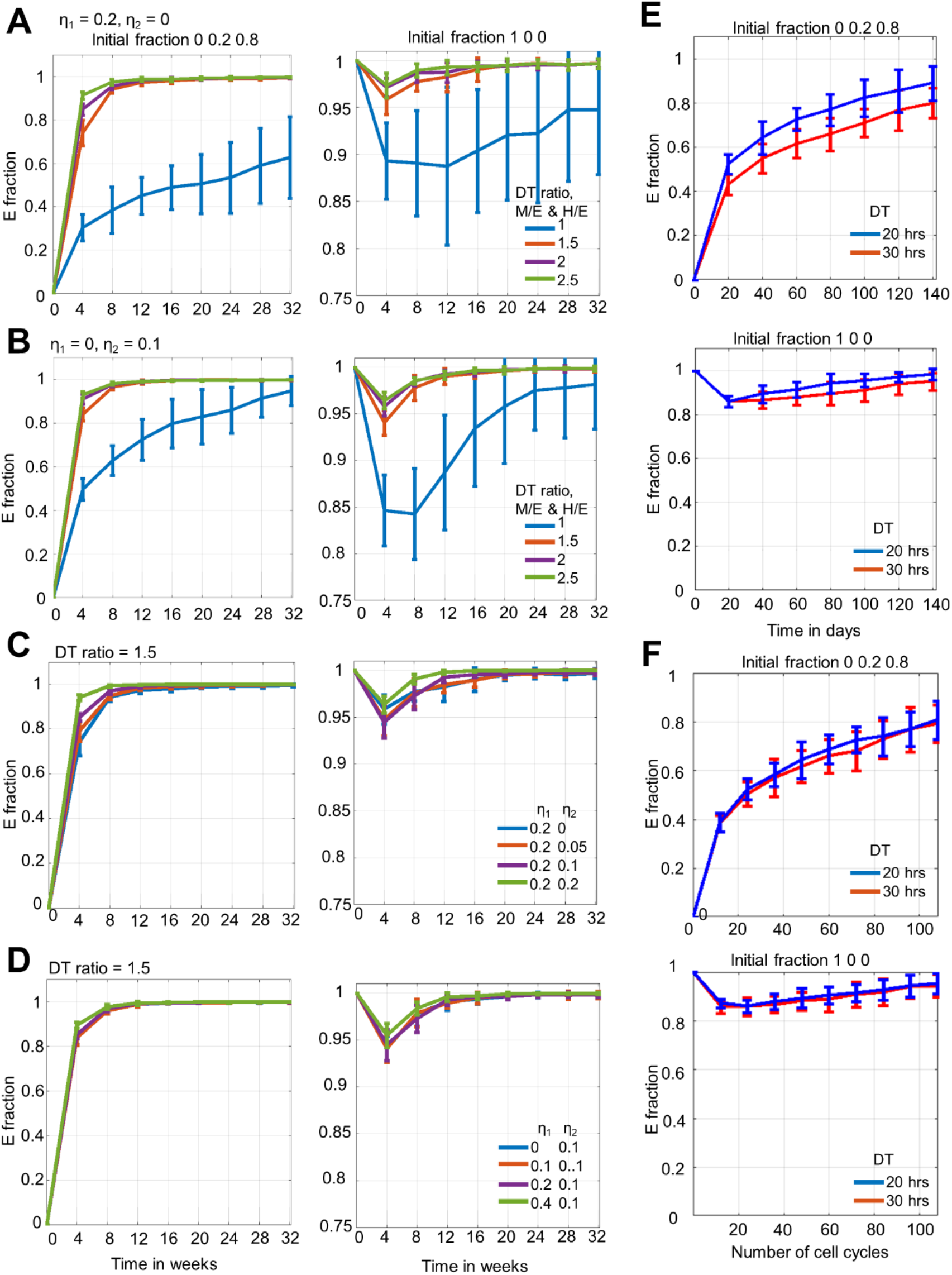
Temporal changes in E fraction for combinations of avg. doubling time (DT) ratio, *η*_1_, and *η*_2_ values; and higher cell doubling time. Temporal E fraction changes for different DT ratios when **(A)** *η*_1_ = 0.2 and *η*_1_ = 0, and **(B)** *η*_1_ = 0 and *η*_1_ = 0.1. Temporal E fraction changes for different *η*_1_ and *η*_2_ values when **(C)** *η*_1_ = 0.2 and *η*_2_ varying, and **(D)** *η*_1_ varying and *η*_2_ = 0.1. Here, the DT(E, E/M, M) = 20 hrs. E). **E)** Overlap in E fraction changes for DT(E, E/M, M) = 20 hrs and DT(E, E/M, M) = 30 hrs when plotted against time in days. **F)** Overlap in E fraction changes for DT(E, E/M, M) = 20 hrs and DT(E, E/M, M) = 30 hrs when plotted against number of cell cycles, determined by DT. In all, initial fractions of 1) Mix of E/M and M, and 2) pure E phenotypes are considered. The initial population size was 200 cells. Mean and standard deviation calculated from 16 independent runs.

**Fig S3.**
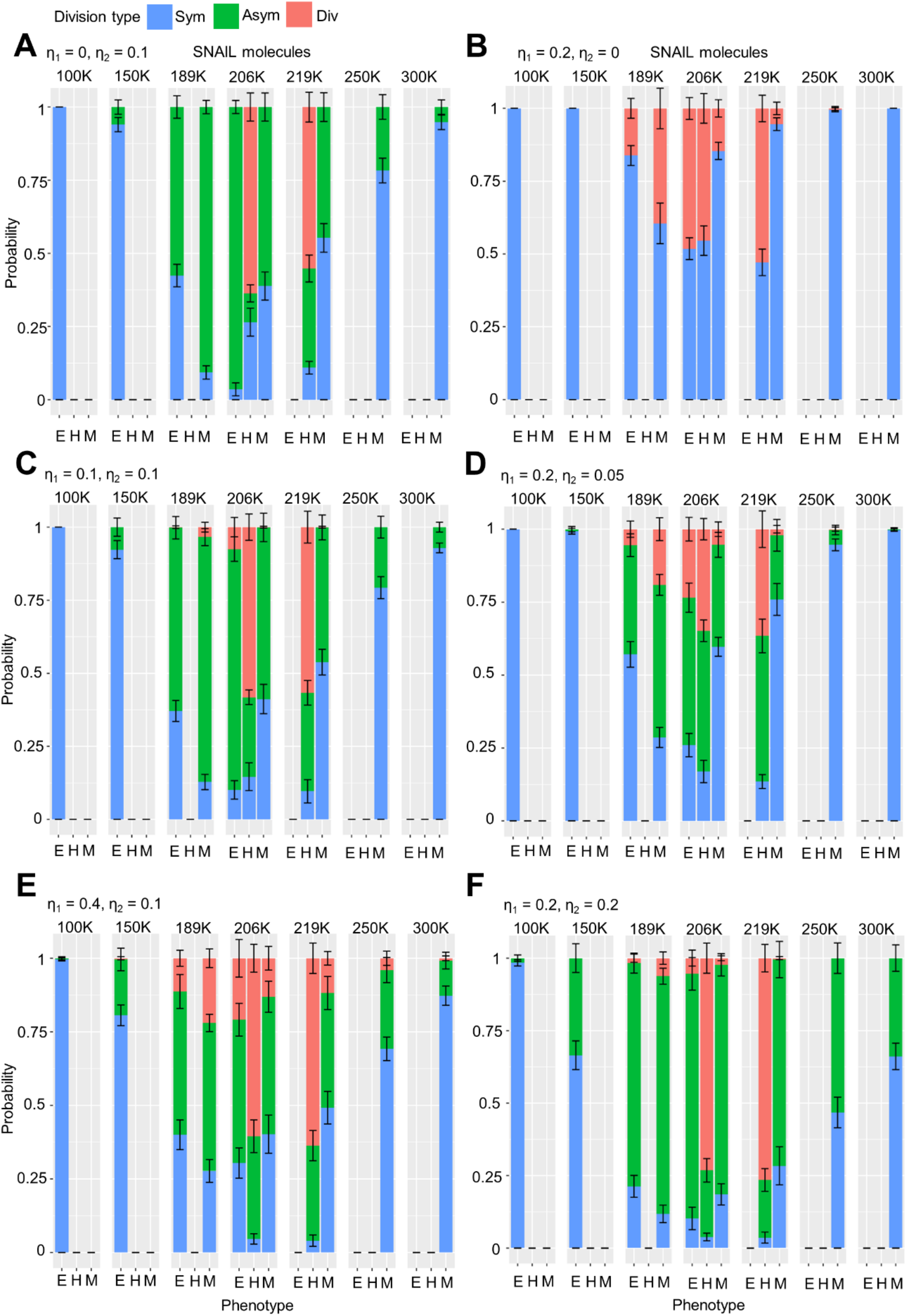
Phenotypic switching probability for various scaling factors (*η*_1_ and *η*_2_) values across SNAIL levels. Mean, and standard deviation are calculated from 10 independent runs of 100 iterations each.

**Fig S4.**
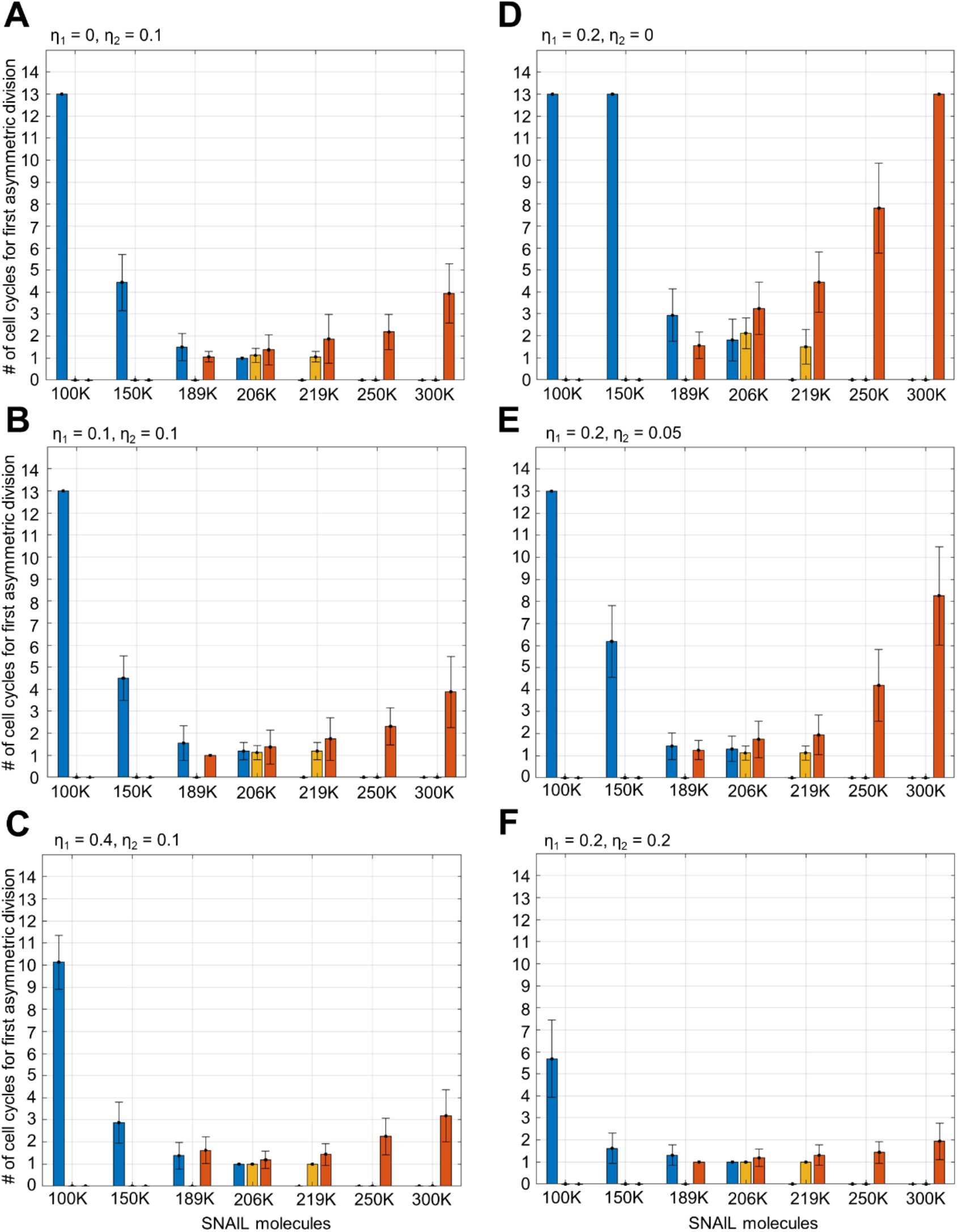
Cell cycles/generations required for first asymmetric switching for various scaling factors (*η*_1_ and *η*_2_) values across SNAIL levels. Progeny up to 12 generations/cell cycles were observed for phenotypic switching. Mean, and standard deviation are calculated from 16 independent runs.

**Fig S5.**
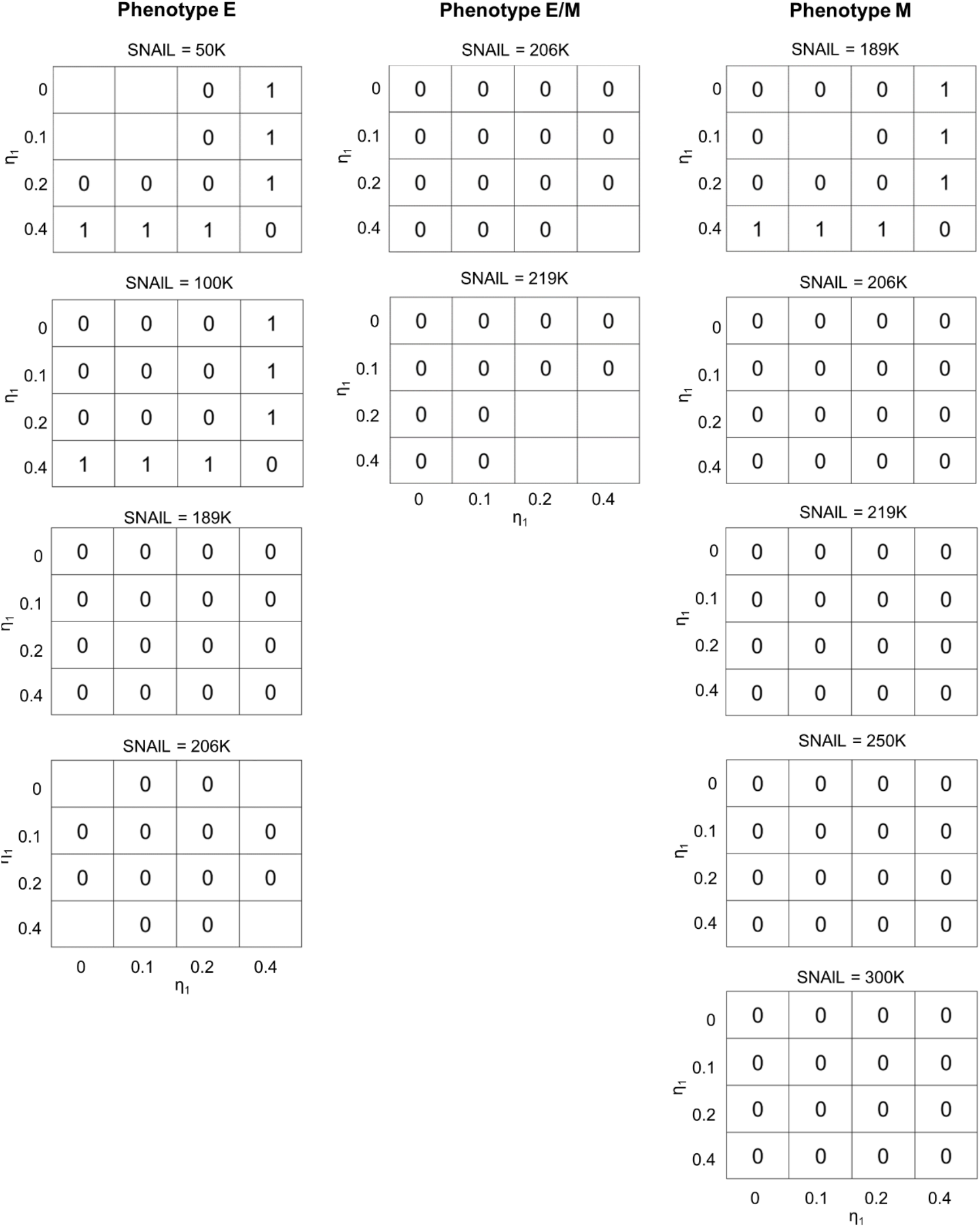
Statistical analysis of differences in minimum cell cycles required for asymmetric division by a cell of given phenotype and SNAIL level on varying *η*_1_ and keeping *η*_2_ fixed (0.1). Here, 1 represents a statistically significant difference, while 0 denotes insignificance, according to p-value of 0.05. The blank space corresponds to nan (not-a-number) values resulted as an outcome of non-variability in data in and across pair of observations.

**Fig S6.**
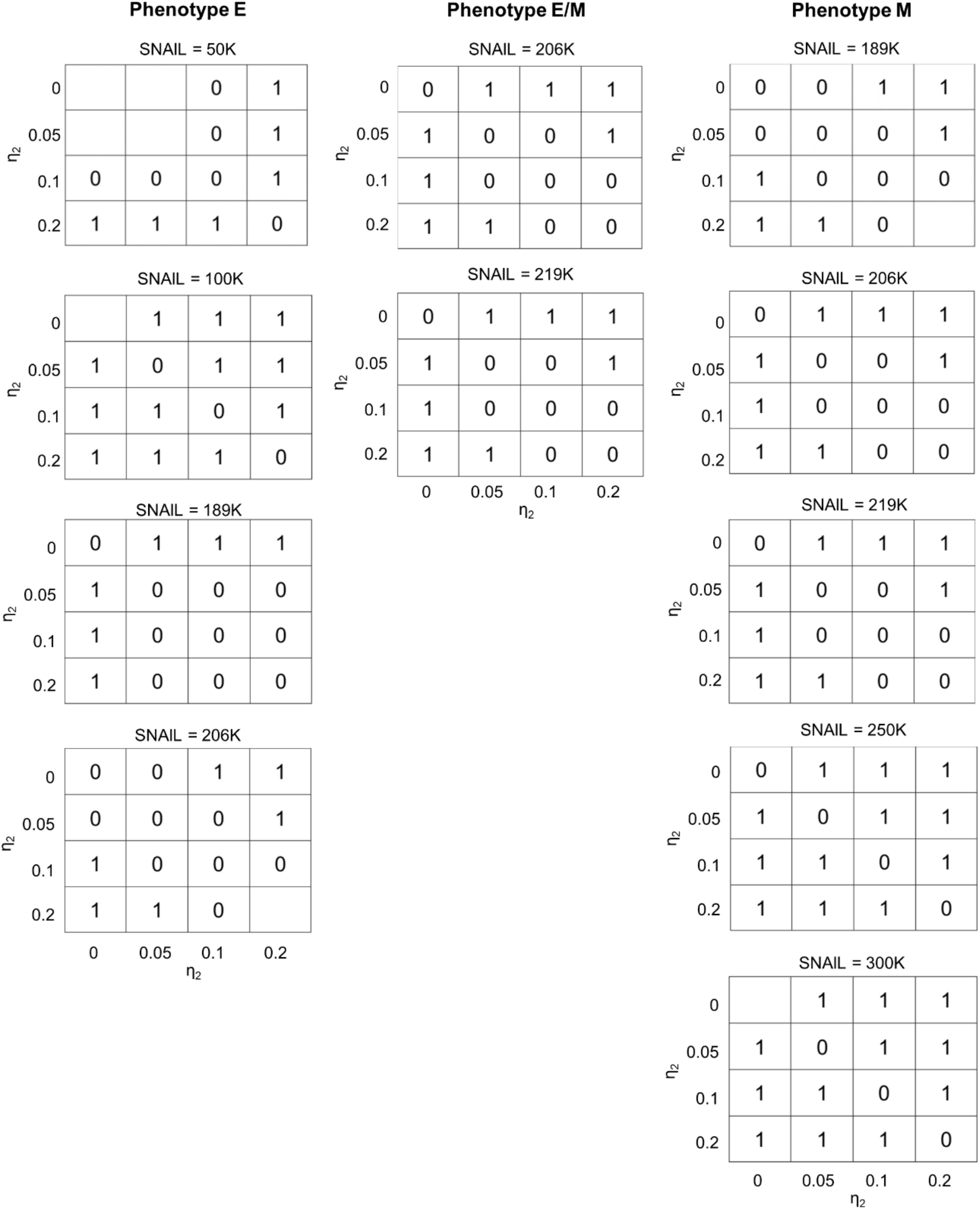
Statistical analysis of differences in minimum cell cycles required for asymmetric division for a cell of given phenotype and SNAIL level on keeping *η*_1_ fixed (0.2) and varying *η*_2_. Here, 1 represents a statistically significant difference, while 0 denotes insignificance, according to p-value of 0.05. The blank space corresponds to nan (not-a-number) values resulted as an outcome of non-variability in data in and across pair of observations.

